# Altered stem cell properties of human hematopoietic stem and progenitor cells based on bone region location

**DOI:** 10.64898/2026.02.25.707977

**Authors:** Christopher J. Wells, Christine Hall, Samantha M. Holmes, Isabelle J. Grenier-Pleau, John F. Rudan, Steve Mann, Sheela A. Abraham

**Affiliations:** Department of Biomedical and Molecular Sciences, Queen’s University, Kingston, Ontario, K7L3N6, Canada; Department of Surgery, Queen’s University, Kingston, Ontario, K7L3N6, Canada

## Abstract

The bone marrow microenvironment forms a highly specialized niche that houses hematopoietic stem and progenitor cells (HSPCs). Within bone, two anatomically distinct regions, the medullary cavity and the trabecular compartment, differ in their cellular and physical composition, with the potential to differentially regulate influence on resident HSPCs. We hypothesized that HSPCs enriched from the medullary cavity (BM) and trabeculae (TB) represent functionally distinct populations. Contrary to this, functional assessment of HSPCs revealed comparable cellular outputs between BM- and TB-derived HSPCs. To investigate whether microenvironmental signaling contributes to functional regulation, we examined the effects of extracellular vesicles (EVs) isolated from medullary BM and TB. Notably, TB-derived EVs inhibited cell cycle progression, directing HSPCs toward a quiescent state. Together, these findings demonstrate that while isolated BM- and TB-derived HSPCs exhibit similar cell-intrinsic properties, EVs enriched from the TB specifically promote HSPC quiescence, supporting a protective regulatory role for the trabecular microenvironment.

## Introduction

The hematopoietic system consists of various cell types and lineages that are governed by the regulation of hematopoietic stem cells (HSCs), that in turn, require lifelong maintenance. These unique cells give rise to differentiated hematopoietic cell populations responsible for immune response, antibody production, oxygen transfer, and blood clotting^1-3^. Because of HSCs innate ability to differentiate, self-renew, migrate, and enter quiescence^4^, this allows the maintenance of clonal populations throughout an individual’s lifetime. It is generally believed that a pool of HSCs are retained over time and remain quiescent, only activated in response to stress and replenishment^5-9^. Despite this, HSPCs are still susceptible to age-related alterations, leading to DNA damage^10^, epigenetic changes^11^, altered colony output, and decreased overall function^12-14^. Therefore, maintaining beneficial signaling and regulation may oppose age-related decline in hematopoietic stem cells and their downstream progenitors.

Stem cell function is intricately linked to their distinct environmental niche. Adult human HSCs reside predominantly within the bone, activating and/or extravasating to the circulation only in response to environmental stressors^15-17^. The bone microenvironment provides a specialized setting, providing key factors conducive to HSC niche formation and overall HSC homeostasis^18,19^. The HSC niche is vital in regulating HSC and HSPC fate by regulating stem cell characteristics through direct cell-cell contact, secreted factors, extracellular matrix, and nutrient access^20,21^. The niche establishes conditions that instruct stem cell behaviour. It is located in proximity to sinusoids with local endothelial cells, perivascular stromal cells, and mesenchymal stromal cells (MSCs)^18,22,23^. The transfer of key factors between these resident niche cells and HSCs is imperative to the controlled regulation of HSCs.

Bone exist as two distinct types based on their corresponding structural and cellular composition, namely compact (cortical) bone and cancellous (spongy) bone. Cortical bone is the dense hard outer layer comprising 80% of total bone mass. The medullary cavity, that houses bone marrow, occupies the space within the diaphysis of long bones and is a hollow chamber surrounded by compact bone, containing vessels and a variety of cell types including MSCs, hematopoietic cells, vascular cells, and adipocytes^24,25^. Within the epiphyses of the long bones sits the trabeculae, a spongy lattice-like structure of bone that comprises bone forming osteoclasts and osteoblasts, HSPCs, lineage-committed precursors, MSCs, and endothelial cells^26,27^. Both the medullary cavity and trabeculae contain hematopoietic bone marrow. Early in development, hematopoietically active red bone marrow dominates the extracellular spaces of the long bones, providing the niche for active and functional HSCs. Aging disrupts the equilibrium, allowing the build-up of fatty deposits in the medullary cavity engendering the formation of yellow bone marrow^28^. As adulthood continues, the quantity of functional red bone marrow decreases, replaced by yellow marrow, pushing the HSPC pool further into specialized microenvironments within the trabeculae^29,30^.

Our group and others have both identified the existence and documented the high concentration of extracellular vesicles (EVs) in human bone. EVs are phospholipid-enclosed nanoparticles (30nm-10microns) containing cell-derived bioactive cargo such as proteins, nucleic acids, lipids, carbohydrates and metabolites^31^. All cells are believed to release EVs, which may facilitate intercellular communication by transferring regulatory factors from donor to recipient cells. Therefore, EVs contribute to the establishment of specific environments by mediating cell to cell interactions and communication^32^. In fact, various cell types harness the mobility of EVs to create favourable local and distant environments that support cellular function^33,34^.

The transfer of multiple factors between resident niche cells and HSCs is well documented^18,35,36^. Extracellular vesicles are established mediators of cellular communication both locally and distantly but remains under researched in the context of human bone. Our group has previously documented the unique role of EVs to define distinct niches between hematopoietic environments, highlighting the potential for EVs to mediate intercellular communication in the bone niche^37^. In addition, our previous work highlighted the specialized role of trabecular EVs to influence HSC function by downregulating cell cycle progression in umbilical cord blood (UCB) HSPCs^37^. From our previous study we began to wonder if HSCs isolated from the distinct bone locations represented unique HSC types or were predominately influenced by niche factors within a specific microenvironment. Results presented here reveal that EVs may play an important role, contributing to maintaining a quiescent population within the trabecular region of the bone.

## Methodology

### Ethics

Human sample collection followed the Declaration of Helsinki and was approved by the Queen’s University Health Sciences and Affiliated Teaching Hospitals Research Ethics Board (HSREB). Approval for the collection of human umbilical cord blood was obtained prior to commencement of the study (Department Code: DBMS-093-18, TRAQ# 6024642). Approval for collection of human blood, bone marrow, and trabeculae was obtained prior to commencement of the study (Department Code: DBMS-093-18, TRAQ# 6036291). Blood donors provided informed verbal consent prior to total hip arthroplasty surgery according to HSREB regulations.

### Human Samples

Human blood, bone marrow, and trabeculae samples were collected from consenting patients undergoing total hip arthroplasty surgery at Hotel Dieu Hospital and Kingston General Hospital in Kingston, ON, Canada. Exclusion criteria included inflammatory arthropathies, cancer, rheumatoid arthritis, any blood disorders, and severe systemic cardiovascular or respiratory disease. Accepted patients were undergoing elective hip replacement and typically diagnosed with osteoarthritis. Samples were transported immediately upon collection in surgery and underwent HSPC and EV enrichment using described protocols.^38^ All samples were processed as outlined in the previously published work.^37,38^

### Enrichment of HSPCs

HSPCs were obtained from umbilical cord blood (UCB) collected in collaboration with Kingston General Hospital, Canada during full-term planned caesarean section surgeries. Immediately upon extraction UCB samples were diluted in citrate dextrose anticoagulant to a final concentration of 25% before transport to the lab. Samples were further diluted 1:1 in PBS and layered onto 20mL of Ficoll-Paque Premium (Sigma-Aldrich). Samples underwent a centrifugation at 300xg (deceleration 0) for 30mins at 21°C and the mononuclear cell (MNC) layer was collected. The MNCs were further purified by positive selection of CD34 expressing HSPCs using the EasySep Human Cord Blood CD34^+^ Selection Kit II (StemCell Technologies).

Bone marrow samples were transported upon collection and diluted in PBS. Samples were centrifuged at 300xg for 10mins at 21C and the cell pellet was separated from the EV-containing biological fluid. They were further washed in PBS supplemented with 2% foetal bovine serum (FBS) and EDTA at a concentration of 1mM (2% FBS/PBS, 1mM EDTA) and centrifuged at 300xg for 7mins at 21°C before layering onto 6mL of Ficoll-Paque Premium (Sigma-Aldrich). Samples underwent a centrifugation at 300xg (deceleration 0) for 30mins at 21°C and the mononuclear cell (MNC) layer was collected before proceeding with EasySep Human Bone Marrow CD34^+^ Selection Kit II (StemCell Technologies) HSPC enrichment.

Trabecular bone samples collected and transported from hospital. Bone was probed and flushed with PBS before rocking 10mins at 4°C. The sample was poured through a 40μm cell strainer and centrifuged at 200xg for 10mins at 21°C. Trabecular cells were separated from EV-containing biological fluid, washed in 2% FBS/PBS 1mM EDTA, and centrifuged at 300xg for 10mins at 21°C before layering on 15mL of Ficoll-Paque Premium (Sigma Aldrich). Samples underwent a centrifugation at 300xg (deceleration 0) for 30mins at 21°C. The MNC layer was collected and underwent positive selection using the EasySep Human Bone Marrow CD34^+^ Selection Kit II (StemCell Technologies). Bone marrow and trabecular HSPCs were either used fresh the same day or frozen in liquid nitrogen (-196°C).

### Enrichment of EVs

Extracellular vesicle-rich supernatant was isolated from the three different hematopoietic environments followed by a 2-step enrichment protocol incorporating iodixanol density cushion (IDC) ultracentrifugation followed by size exclusion chromatography outlined in our paper^38^. Whole blood was collected in EDTA K2 tubes and centrifuged at 1900xg (deceleration 0) for 10mins at 21°C, followed by a second centrifugation at 2500xg for 10mins at 21°C. Bone marrow samples were transferred from the hospital in EDTA K2 tubes and washed with PBS. Bone marrow samples were centrifuged at 300g for 10mins at 21°C to collect cell-free biological fluid. Bone marrow biological fluid was washed once more before being concentrated using an Amicon Ultra-15 centrifuge filter (10kDa) (Miltenyi Biotech) to a final concentration of 6mL. Trabecular bone samples were probed and flushed with PBS before rocking 10mins at 4°C. The sample was poured through a 40μm cell strainer and centrifuged at 200xg for 10mins at 21°C. After removal of the fat layer, trabecular cell-free biological fluid was collected and concentrated using an Amicon Ultra-15 centrifuge filters (10kDa) (Miltenyi Bitotech) to a final volume of 6mL.

Samples were centrifuged at 2500xg for 10mins at 21°C and strained through a 40μm cell strainer to remove any remaining fat and cell debris. Plasma, bone marrow, and trabeculae samples were layered onto an iodixanol density cushion of Optiprep (Cedarlane) with layers at 50%, 30%, and 10% and placed in an SW41Ti ultracentrifuge rotor. Samples were centrifuged at 178,000xg for 2hrs at 4°C using an LS-60M Ultracentrifuge (Beckman Coulter, USA, cat no. 347240). The high-density layers (interface between 30% and 10% layers) were collected and placed onto SEC columns. Twenty fractions of 0.5mL were collected and EV-containing fractions (F7-12) were pooled and aliquoted. Samples were stored at -80°C. EV sample preparation and enrichment followed the protocols outlined^38^. An aliquot of each EV sample was thawed on ice and characterized using nanoparticle tracking analysis (NTA) and Qubit Protein Assay Kit (ThermoFisher Scientific) to determine particle concentration, size, and protein concentration.

### Estimated Trabeculae Volume Calculation

Trabecular bone weight was determined by weighing the bone on the day of the surgery as well as weighing after 7-10 days of drying. After drying, bone was rehydrated in ddH_2_0 for 24 hours before weighing. Available space is then calculated to provide trabeculae sample volumes. All trabecular bone calculations were conducted as outlined in the paper explaining trabeculae extracellular vesicle enrichment and bone available space calculations.^38^ Bone calculation has been verified and is consistent across individuals^37^. Samples used sex-specific fractions of available trabecular bone space; females = 0.258 and males = 0.203.

### Super Resolution Microscopy of EVs

EVs were captured using the ONi human EV profiler kit v2.0 and processed using stochastic optical reconstruction microscopy (STORM) on the ONi super-resolution nanoimager. Samples were immobilized by phosphatidyl serine (PS) capture on microfluidic glass slides before washing of free capture antibodies. EVs were fixed and stained with a 3-colour tetraspanin antibody panel (CD9-488nm, CD63-561nm, CD81-647nm) using 50%, 40%, and 30% power on 488, 561, and 640 nm lasers, respectively. EVs were quantified based on positive expression of tetraspanin markers and labeled as single, double, or triple positive. EVs were classified as positive for a marker when 10 or more individual localizations were detected in the same channel at a radius of 100nm around the centre of a cluster.

### HSPC Incubation of EVs

CD34+ HSPCs were incubated in serum-free media (SFM), prepared using Iscove’s Modified Dulbecco’s Medium (Sigma-Aldrich), bovine serum albumin, insulin, transferrin (B.I.T.) serum substitute, β-mercaptoethanol (ThermoFisher Scientific), low-density lipoprotein (Sigma-Aldrich), L-Glutamine (ThermoFisher Scientific), and penicillin/streptomycin (ThermoFisher Scientific). Media was further supplemented with cytokines and growth factors IL-3 (20ng/mL), IL-6 (20ng/mL), G-CSF (20ng/mL), Flt3 (100ng/mL), and SCF (100ng/mL). HSPCs were plated at 200,000 cells/mL and incubated for 48hrs in SFM containing EVs at 10μg/mL of EV protein or PBS control. To validate functionality within and between individuals, HSPC and EV incubation experiments included both HSPCs that received EVs from paired source and HSPCs that received EVs from different individuals. HSPC source (BM and TB) and EV source (BM and TB) were always kept consistent throughout comparisons. For all incubations, cells were incubated at 37°C with 5% CO_2_.

### Flow Cytometric Analyses

Multiparameter flow cytometry was used for immunophenotyping^39-41^ and functional analyses. HSPC cell populations and lineages were identified by immunophenotyping using monoclonal antibodies to human CD34 (581), CD38 (HIT2), CD90 (5E10), CD45RA (HI100), CD123 (6H6), CD10 (HI10a), and CD49f (G0H3) from BD Biosciences. Cells were stained in 2% FBS/PBS, 1mM EDTA solution for 45mins at 4°C and were washed twice prior to analysis. Cell cycle status was assessed by Ki67 and DAPI staining. Cells were first fixed with 1.5% PFA for 30mins at 4°C and then permeabilized with ice-cold 100% acetone for 10mins at 4°C. Cells were stained with Ki-67 for 30mins at 21°C prior to staining with CD34 for 45mins at 4°C. Cells were washed once and then incubated with DAPI on ice for 7mins before a second wash. Proliferation kinetics of HSPCs was performed by staining with carboxyfluorescein succinimidyl ester (CFSE) fluorescent dye. CD34^+^ HSPCs were stained with CFSE for 10mins at 37°C before a 5-day incubation in SFM supplemented with cytokines and growth factors. On Day 5, cells were stained with CD34 for 45mins at 4°C and then assessed for cellular divisions across the 5-day incubation. For experiments assessing HSPCs only, cells were used fresh after isolation or were thawed and allowed to rest overnight before analyses. Experiments assessing HSPCs incubated with EVs, HSPCs were thawed and allowed to rest overnight before undergoing a 48hr incubation with 10μg/mL of EV protein as outlined above. All experiments included isotype controls or Fluorescent-Minus-One (FMO) controls using primary cells. Flow cytometric experiments were performed on FACSAria III (BD Biosciences), FACSymphony A3 (BD Biosciences, or a CytoFLEX S (Beckman Coulter) and analyzed using FlowJo v10.10.0 software.

### Colony-Forming Cell (CFC) Assay

Colony formation capacity of HSPCs alone was evaluated using freshly isolated CD34^+^ cells. 1000 HSPCs were plated into 1.2mL of H4435-enriched Methocult Medium (StemCell Technologies) for each condition. Conditions were plated in triplicate into 35mm dishes and incubated for 10-12 days at 37°C with 5% CO_2_.

Bone EV assessment incubated HSPCs with 10ug/mL of EV protein for 48hr as outlined above. Conditions were counted and 1000 cells were plated in triplicate into 1.2mL of H4435-enriched Methocult Medium (StemCell Technologies). Conditions were plated into 35mm dishes and incubated for 10-12 days at 37°C with 5% CO_2_. For both experiments, colonies were quantified and qualified based on morphology using a light microscope. Colony types include colony-forming unit granulocyte/erythrocyte/macrophage/megakaryocyte (CFU-GEMM), colony-forming unit-granulocyte/macrophage (CFU-GM), colony-forming unit-granulocyte (CFU-G), colony-forming unit-macrophage (CFU-M), colony-forming unit-erythroid (CFU-E), and burst-forming unit-erythroid (BFU-E).

### Western Blot Analysis

EV samples were diluted 1:10 with 10X RIPA and incubated for 30 mins at 4°C. Due to increased surface area for lysis, tubes were agitated every 5mins throughout the 30mins incubation. Protein lysates were centrifuged at 13,000 rpm for 5 mins. Samples were diluted 1:4 in 4X sample buffer (ddH2O included if further dilution required) and heated at 95°C for 5 mins before proceeding to Western blot analysis. Western blot gels were created using 1.5 M Tris-Cl (pH=8.8) base resolving gel and 0.5 M Tris-Cl (pH=6.8) base stacking gel. Each channel was loaded with 20 ug of EV protein lysates and run alongside 8-260 kDa protein ladder. Gels were transferred onto PVDF membranes and blocked for 1 hr. Sample membranes were incubated 1:500 or 1:1000 with primary antibodies (CD63, CD9, SDCBP) at 4°C overnight. Membranes are washed of primary antibodies using TBS-T. Secondary antibodies are then incubated 1:5000 using Dylight 680 or Dylight 800 (Invitrogen, USA) at room temperature for 1 hr. Membranes were washed and then imaged on Odyssey CLx infrared imaging system.

## Results

### Bone marrow and trabeculae HSPCs exhibit similar function, and both contribute to the HSPC pool within the bone

The rarity and difficulty of obtaining human hematopoietic stem and progenitor cell populations limits fine analyses of the distinct pools within the hematopoietic system. Current experiments rely overwhelmingly on human HSPCs derived from either bone marrow aspirates (BM). As a first assessment, HSPC concentration was calculated between the BM and TB (Fig. 1B). The percentage of HSPCs, determined by positive expression of the surface marker CD34, remained consistent between both environments (Fig. 1B). HSPCs from the 2 different sources were then further characterized using flow cytometric immunophenotyping to identify specific cell populations within the hematopoietic hierarchy (Fig. 1C). CD34 and CD38 expression allows identification of the broad populations of HSPCs. HSPCs from BM and TB showed no differences in the specific primitive populations, the long-term HSCs (LT-HSCs; CD34^+^,CD38^-^,CD90^+^,CD45RA^-^,CD49f^+^), short-term HSCs (ST-HSCs; CD34^+^,CD38^-^,CD90^-^,CD45RA^-^,CD49f^+^), 90+ HSCs (CD34^+^,CD38^-^,CD90^+^,CD45RA^-^,CD49f^-^), and 90-HSCs (CD34^+^,CD38^-^,CD90^-^,CD45RA^-^,CD49f^-^) (Fig. 1D). Similarly, the progenitor populations show no differences between HSPCs of the BM and TB pools; megakaryocyte-erythroid progenitor (MEP; CD34^+^,CD38^+^,CD10^-^,CD45RA^-^,CD123^-^), common myeloid progenitor (CMP; CD34^+^,CD38^+^,CD10^-^, CD45RA^-^,CD123^+^), granulocyte-macrophage progenitor (GMP; CD34^+^,38^+^,CD10^-^, CD45RA^+^,CD123^+^), lymphoid-primed multipotent progenitor (LMPP; CD34^+^,CD38^-^, CD45RA^+^,CD90^-^,CD10^-^), and NK-B cell progenitor (NK/B Prog; CD34^+^,CD38^-^, CD45RA^-^,CD90^-^,CD10^+^) (Fig. 1E-F). The differences seen in the CD34^+^38^-^ populations is driven by the decrease in the lymphoid lineage (multiple lymphoid progenitors (MLPs; CD34^+^,CD38^+^,CD45RA^+^,CD10^+^)) within TB HSPCs. These findings suggest slight variations in the BM- and TB-derived HSPCs pools driven by shifts in subpopulations of the hematopoietic hierarchy.

**Figure 1.**
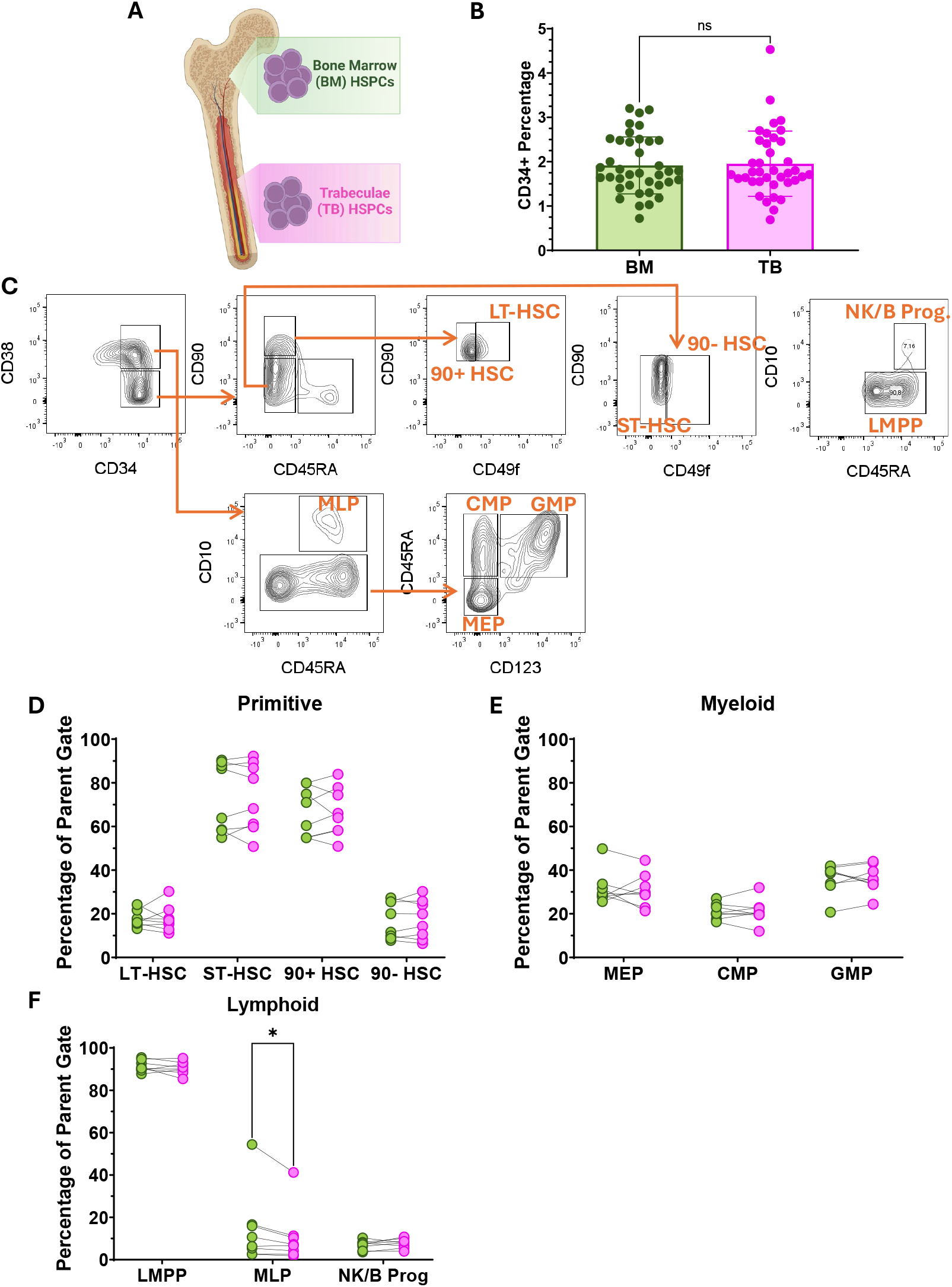
HSPC characterization of CD34+ cells isolated from paired bone marrow (BM) and trabeculae (TB). Characterization of paired HSPCs isolated from the bone marrow (BM) and trabeculae (TB) of individuals undergoing total hip arthroplasty surgery. (**a**) Graphic highlighting different origins of bone marrow (BM) and trabeculae (TB) derived HSPCs. (**b**) Percentage of CD34+ cells within the mononuclear (MNC) cell population of isolated samples (n = 38) assessed by paired t-test. (**c**) Flow cytometric gating strategy to identify HSPC subpopulations after singlet gating. (**d-f**) Flow cytometry HSPC panel of paired HSPC samples indicated connected points (n = 8). Comparison of HSPC subpopulations (**d**) stem cell populations (**e**) myeloid progenitors (**f**) lymphoid progenitors. A two-way repeated measures ANOVA with Šidák’s multiple comparisons test was computed for each comparison * *P* ≤ 0.05, ** *P* ≤ 0.01, *** *P* ≤ 0.001, **** *P* ≤ 0.0001

Despite potential differences in the HSPC pool between the two bone environments, cells must also be assessed for key stem cell characteristics to validate HSPC functionality. First, HSPCs were assessed on their ability to form myeloid colonies via a colony cell forming (CFC) assay. A traditional assay used to assess the potential for multipotency of HSPCs, this experiment evaluates the production of myeloid and erythroid colonies across 10-day incubations. Literature identifies, BM HSPCs demonstrate significant reductions in total colon formation compared to UCB HSPCs after 10 days^42,43^. This observation appears to be consistent across all bone derived HSPCs, with TB HSPCs also producing similar total colonies to BM HPSCs (Fig. 2A).

**Figure 2.**
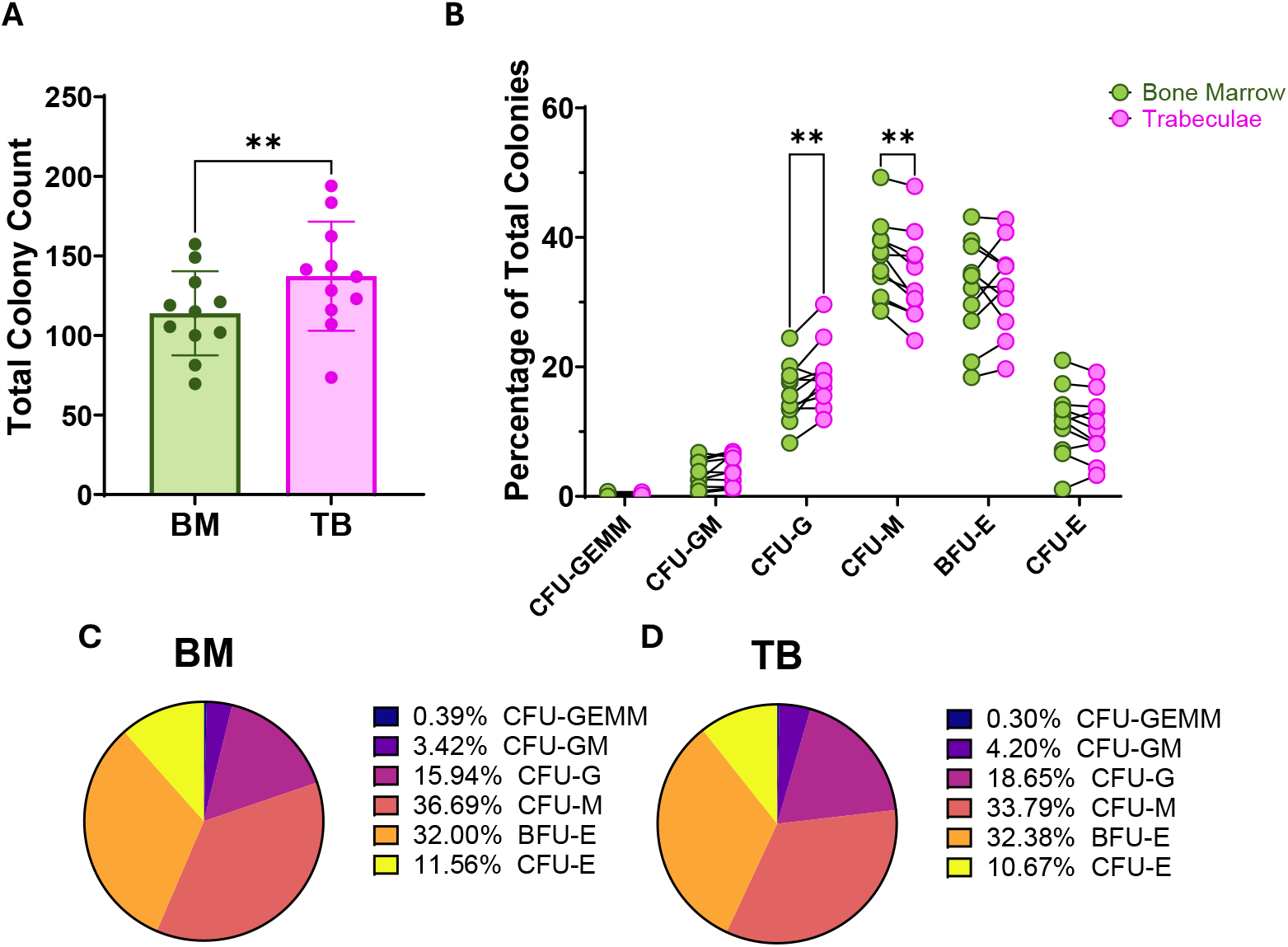
Functional Analysis of bone marrow and trabeculae CD34+ HSPCs colony forming capacity. Functional comparisons of HSPCs isolated from paired BM and TB samples (n = 11). (**a**) Total colony counts after 10-day incubation in methylcellulose analyzed by Paired t-test. (**b**) Colony breakdown as percentage of total colonies with paired samples connected; colony-forming unit-granulocyte/erythroid/macrophage/megakaryocyte (CFU-GEMM), colony-forming unit-granulocyte/macrophage (CFU-GM), colony-forming unit-granulocyte (CFU-G), colony-forming unit-macrophage (CFU-M), burst-forming unit-erythroid (BFU-E), and colony-forming unit-erythroid (CFU-E). A Two-way repeated measures ANOVA with Šidák’s multiple comparisons test was computed for each comparison. Pie chart comparison of colony types averaged across individuals between (**c**) BM, and (**d**) TB HSPCs. * *P* ≤ 0.05, ** *P* ≤ 0.01, *** *P* ≤ 0.001, **** *P* ≤ 0.0001

Although, TB HSPCs do produce more total colonies than BM HSPCs isolated from the same individual, suggesting a potential difference in colony formation ability between HSPCs of different environments (Fig. 2A). When partitioning total colonies into specific colony lineages/types by proportion, both bone-derived HSPCs exhibit similar production of all colony lineages, implying the total colony count differences are regardless of lineage. However, there is a shift in TB HSPCs away from macrophage colonies (CFU-M) towards granulocyte colonies (CFU-G) compared to BM HSPCs (Fig. 2B). Population differences are further highlighted upon visualization by pie chart, demonstrating comparable averaged proportions of colony types across individuals (Fig. 2C-D). These findings demonstrate that BM- and TB-derived HSPCs express similar colony formation. Further evaluation of multipotent potential is required to confirm the basis for differences in total colony production between BM- and TB-derived HSPCs.

Colony formation by the HSPC pool could be a result of either increased proliferative potential leading to higher total colony units or higher multipotency activated by exiting from quiescence. To determine whether the differences in colony formation are due to proliferation or quiescent capabilities; BM- and TB-derived HSPCs underwent flow cytometric analysis. Shifts in cell cycle transitions between HSPCs sourced from BM vs TB were ascertained by staining with Ki-67 and DAPI then analyzed by flow cytometry (Fig. 3A). State specific data suggests that bone HSPCs demonstrate similar cell cycle dynamics regardless of environment between G0, G1, S, and G2/M phases (Fig. 3B-E). Identification of the cell cycle status confirms that colony formation was not due to differences in the quiescent populations (G0) after analyzing cell cycle downstream of cell isolation (Fig. 3B). Proliferation potential was also monitored over 5 days and quantified by carboxyfluorescein succinimidyl ester (CFSE) fluorescence. CD34^+^ HSPCs were stained with CFSE and incubated at 37°C and 5% CO_2_ for 5 days (Fig. 3F). On day 5, cellular divisions were quantified across the entire cell population (Fig. 3G), as well as the cells that retained CD34 expression (Fig. 3H). BM and TB HSPCs demonstrate similar proliferation potential across the entire cell population, both by location and within individuals (Fig. 3G). In the cell population that retained CD34 expression, BM HSPCs contained a greater percentage of undivided cells (D0). However, the two HSPC populations showed similar proportions of cells in all consecutive divisions, indicating comparable proliferation dynamics (Fig. 3H). The combined findings suggest there are no differences between BM- and TB-sourced HSPC function.

**Figure 3.**
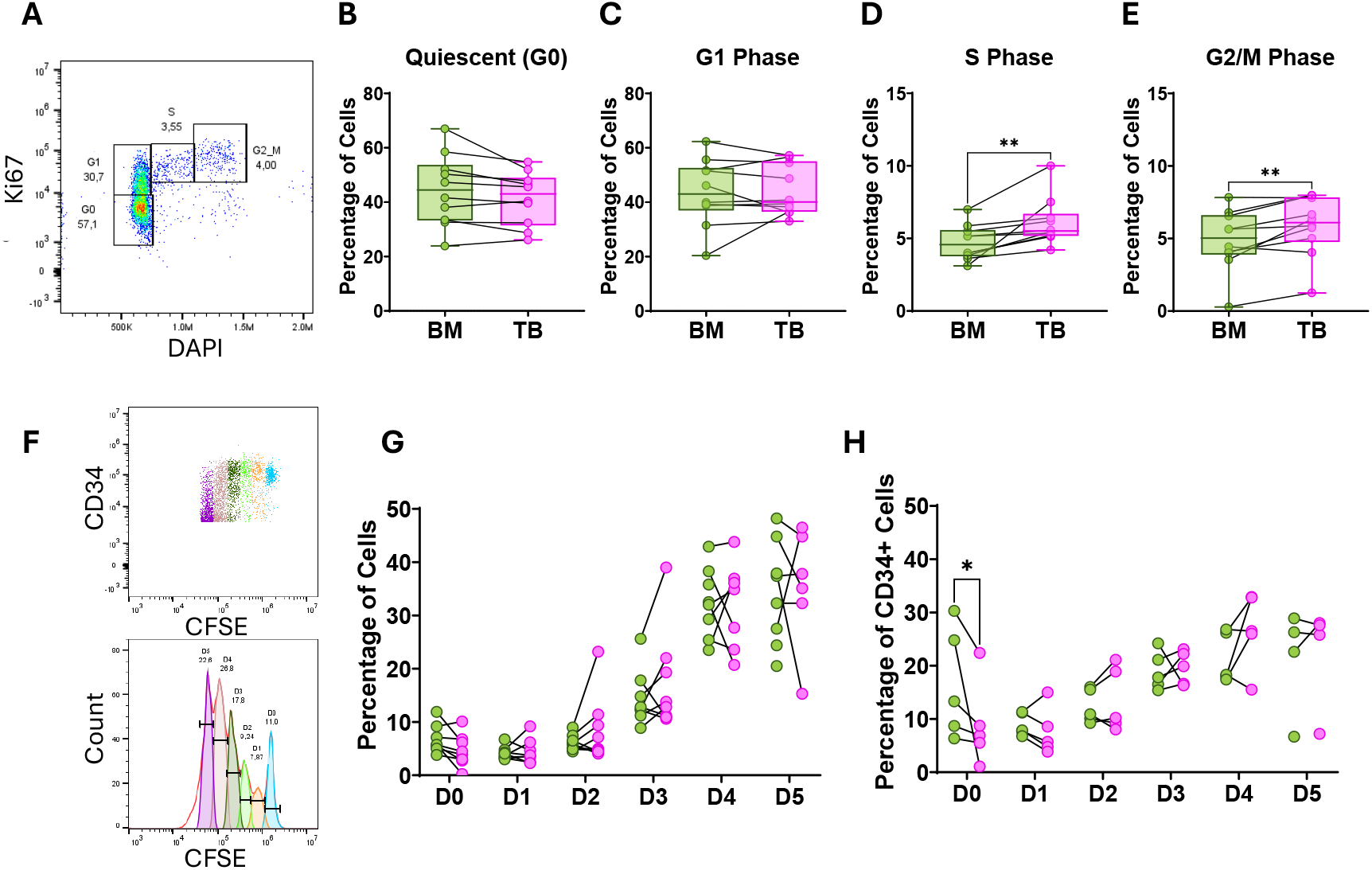
Cell cycle dynamics and proliferative comparison of BM and TB derived HSPCs. (**a**) Representative flow cytometry gating of cell cycle populations. Breakdown of BM (n=10), and TB (n=10) HSPCs indicating paired samples separated into cell cycle phases (**b**) quiescent (G0) populations (**c**) G1 phase, (**d**) S phase, and (**e**) G2/M phases. A paired t-test was computed for each cell cycle phase comparison. (**f**) Representative flow cytometry gating of CFSE labeled cellular divisions. (**g**) Cellular divisions of paired BM and TB HSPCs (n=8) over 5-day incubation. (**h**) Percentage of cells undergoing cellular divisions while maintaining CD34 surface marker after 5-day incubation. CFSE comparisons computed by mixed-effects analysis with Šidák’s multiple comparisons tests. * *P* ≤ 0.05, ** *P* ≤ 0.01, *** *P* ≤ 0.001, **** *P* ≤ 0.0001

### Extracellular vesicles from different bone locations exert differential effects on HSPCs

In our previously published work, TB EVs were able to promote quiescence through downregulation of cell cycle progression in umbilical cord sourced HSCs.^37^ In this study, EV enrichment was conducted across all individuals therefor directly comparing BM and TB HSPC from the same individual (Fig. 4A). Samples underwent Western blot validation of common EV markers (CD63, CD81, SDCBP (Syntenin)) expressed on EVs (Fig. 4B). EVs were then characterized based on size and concentration, remaining consistent with previous publications observing significantly greater concentrations of particles in the BM than in the TB (Fig. 4C). As well, TB EVs were significantly larger than EVs isolated from the bone marrow of the same individual but similar average protein concentration (Fig. 4D-E). Super resolution microscopy was then conducted to visualize individual particles. Bone derived EV samples were assessed for tetraspanin expression of CD63 (yellow), CD81 (pink), and CD9 (blue) (Fig. 4F). Individual particles were classified as single, double, or triple positive based on localizations of fluorescence and quantified as percentage of total population (Fig. 4G).

**Figure 4.**
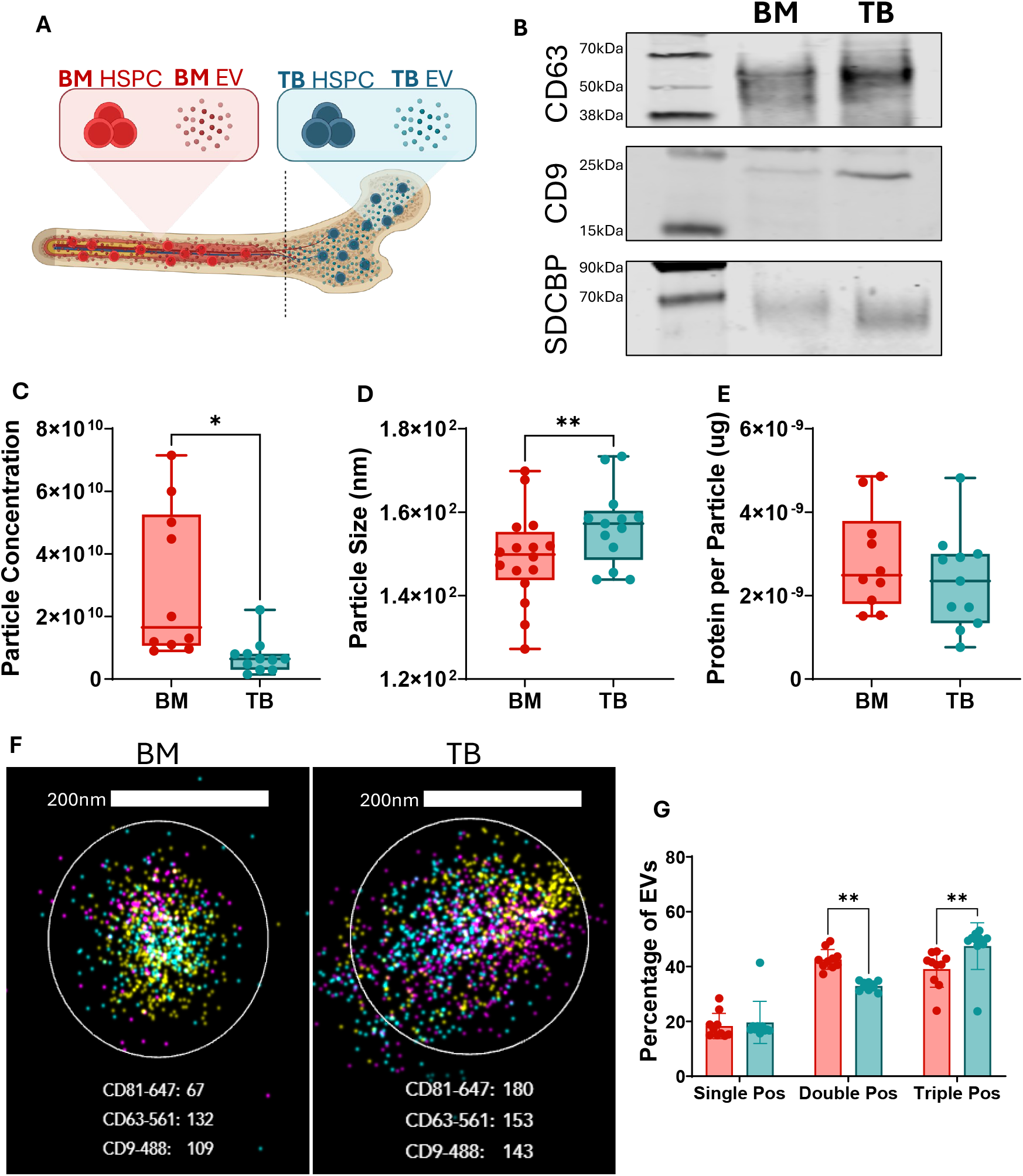
Bone marrow (BM) and trabeculae (TB) derived extracellular vesicle characterization. (**a**). Graphic depicting bone marrow and trabeculae derived HSPCs and EVs. (**b**) Western blot confirmation of EV markers CD63, CD9, and SDCBP (Syntenin) expression in BM and TB EVs. Extracellular vesicles enriched rom bone marrow (TB), and trabeculae (TB) characterized by (**c**) size, (**d**) particle concentration, and (**e**) average protein per particle (n=12). (**f**) Super resolution microscopy images of EVs isolated from TB, and BM labeled with CD63 (yellow), CD81 (pink), and CD9 (blue). (**g**) Individual particle identification of tetraspanin (CD63, CD81, CD9) expression characterized as single, double, or tripled positive (n=1). * *P* ≤ 0.05, ** *P* ≤ 0.01, *** *P* ≤ 0.001, **** *P* ≤ 0.0001

To validate if EVs impart consistent functional effects to BM- and TB-derived HSPCs as to UCB HSPCs, in vitro assays were conducted incubating HSPCs with EVs over 48hrs (Fig. 5A). Colony-forming cell assays were conducted to compare the colony formation of BM- or TB-derived HSPCs treated either with BM EVs, TB EVs, or PBS controls. Cells were incubated with 10μg/mL of EV protein for 48 hrs, followed by plating into methylcellulose media supplemented with HSPC specific growth factors. Corresponding colony counts validate previous findings that TB EVs cause decreased total colony accounts in HSPCs regardless of environmental origin (Fig. 5B). To account for biological variability between individuals, colony totals were standardized to the PBS control of the same cell source, further highlighting the significant reductive effect of TB EV-treatment on HSPCs (Fig. 5B). Colony lineage formation exhibits similar profiles between for both HSPC populations across EV treatments, in all except the most differentiated CFU-E colonies decreasing in the TB treated with TB EVs (Fig. 5B-C). The findings confirm TB EVs reduce total colony count in all HSPCs and necessitates the question of whether TB EVs also alter cell cycle transitions in bone HSPCs despite age-associated differences to UCB HSPCs. Therefore, interrogation of cell cycle transitions was conducted to observe potential differences in quiescent (G0) populations in response to BM and TB EV stimulation in both BM- and TB-derived HSPCs. In BM-derived HSPCs, TB EV incubation significantly promoted cellular quiescence compared to BM EV treatment (Fig. 5D). When evaluating TB EV treatment in BM HSPCs, only G0 was significantly altered, whereas there were slight changes to populations in G1, S, and G2/M phases (Fig. 5E). This trend continued when observing EV treatment in TB-derived HSPCs, highlighting a significant increase in the quiescent HSPC population with treatment of TB EVs (Fig. 5 F). As well, no shifts were seen in the other cell cycle phases for treated TB HSPCs (Fig. 5G). Overall, this data further validates that TB EVs target HSPCs, regardless of origin, to downregulate colony formation and promote entry into quiescence. Therefore, the microenvironment within the trabeculae may provide a protective environment for HSPCs and specifically foster HSC regulation via a nano-based niche controlled by EVs.

**Figure 5.**
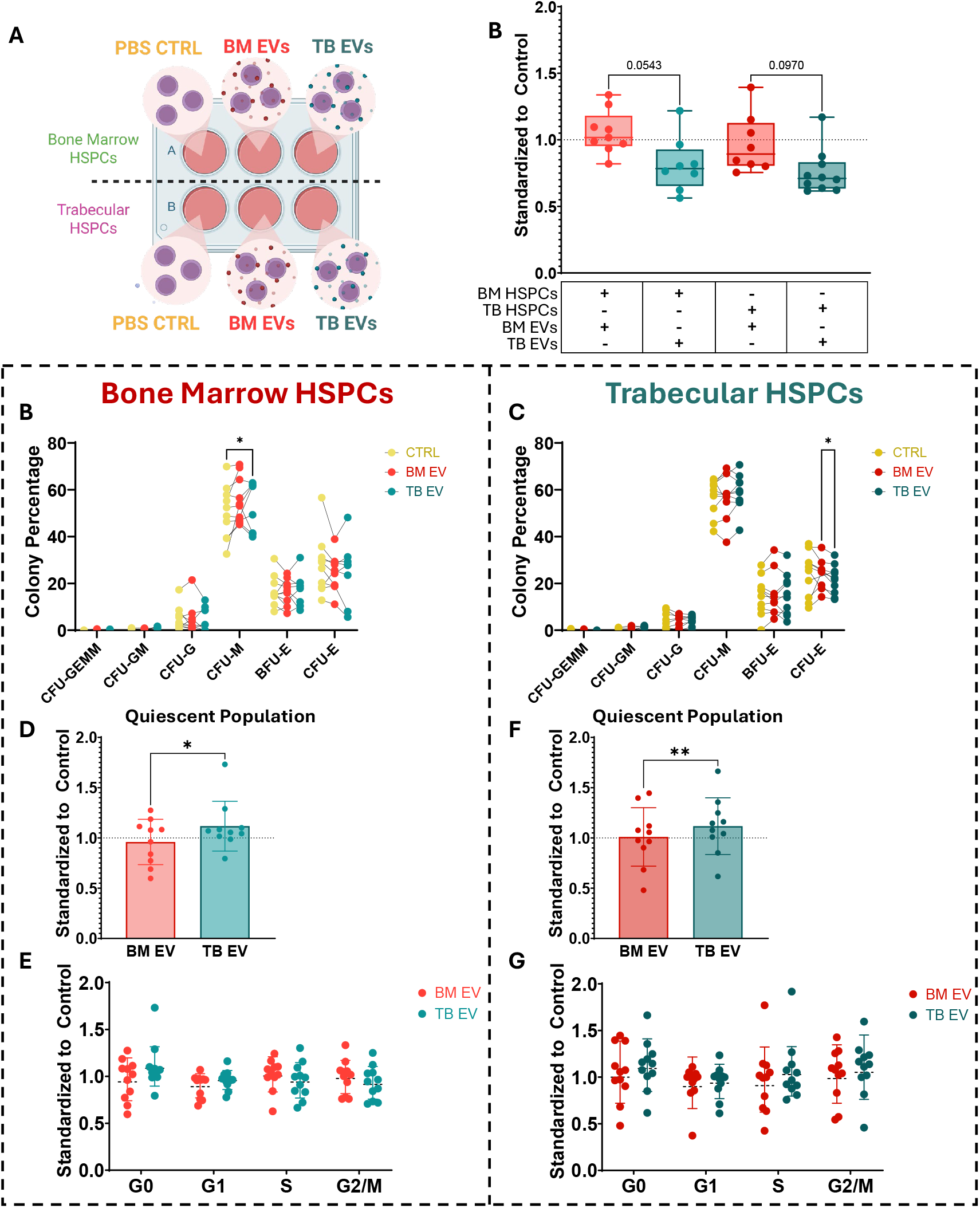
Comparison of bone EV-treatment on bone marrow and trabeculae HSPCs. (**a**) Incubation outline for BM and TB HSPCs treated with PBS control, BM EVs, or TB EVs. (**b**) Total colonies standardized to PBS control of paired BM and TB HSPCs samples treated with BM EVs (red) or TB EVs (blue). Colony type breakdown of (**c**) BM treated and (**d**) TB treated HSPCs. Cell cycle analysis by Ki67 and DAPI staining of BM HSPCs (e-f) and TB HSPCs (g-h) standardized to PBS control. (**e**) Quiescent population comparison. (**f**) Sample separated into cell cycle transition states. (**g**) Quiescent population of TB HSPCs. (**h**) TB samples separated into cell cycle transition states. * *P* ≤ 0.05, ** *P* ≤ 0.01, *** *P* ≤ 0.001, **** *P* ≤ 0.0001

## Discussion

These findings outlined above represent the first study directly comparing HSPCs within differing human bone locations, highlighting the importance of EV based mechanisms that contribute retaining HSPCs in quiescence. The majority of adult HSPC research has centered on cells isolated and impacted by the bone marrow microenvironment^44,45^, despite the trabeculae housing the more haematopoietically productive red marrow^46-48^. Therefore, the trabeculae represent an understudied environment impacting HSPC functionality. This research has sourced two hematological environments from patients undergoing total hip arthroplasty, to determine differences in the resident HSPC population and the impact of its surrounding environment. We have shown that the HSPC pool in the bone marrow and trabecular are similar in functionality and instead impacted by their extracellular vesicle environment. This builds on previously published research by this lab, that identified a new function of trabecular EVs, that provide a protective environment for primitive hematopoietic stem cells^37^.

Hematopoietic stem cell processes have been investigated over the many decades of groundwork^49-54^. Over the years, this has expanded from understanding the function of these rare cells, to the difference in specific populations located within the human anatomy. The data presented here, validates the trabeculae as an attainable source of HSPCs that demonstrate analogous hematopoietic ability to HSPCs from the bone marrow^55^. HSPCs derived from the bone marrow and trabeculae of a single individual represent functionally comparable cell populations, that that can be separated into specific cell types and demonstrate consistent colony formation lineages. It is important to highlight that this data applies to samples within the same individual as well as similarities between environments of different individuals. The findings shown here provide broad characterization of the differences in the HSPCs of the bone marrow and trabeculae, and with more intricate studies, could look at disparities in the most subtle HSPC populations within these environments.

With regards to the environmental regulation of HSPCs, EVs represent another factor contributing to the maintenance of HSPCs and remain source-specific across individuals.

The importance of HSPCs maintaining stem cell functionality to safeguard against aging-associated pathologies^13,56-58^, requires understanding the biology contributing protective environments for HSPCs^59,60^. Our results identify trabecular EVs as an important modulator of stem cell characteristics, driven by their environmental origin. Trabecular EVs provide another avenue into understanding the microenvironment optimal for HSPC maintenance highlighted by their ability to drive change regardless of HSPC origin.

This research demonstrates functional analysis of hematopoietic stem and progenitor cells isolated from a previously under researched hematopoietic environment. We demonstrate that HSPCs isolated from trabecular bone contribute to the HSPC pool and functionally represent an extension of the cell population found within the bone marrow^61^. However, we propose that HSPCs are differentially regulated by the environment in which they reside, partitioning the bone marrow and trabeculae into two unique HSPC pools based on stimulation from their microenvironment that include extracellular vesicles, as well as other external cellular factors.

The role of EVs in cellular communication, and the debate whether EVs are a directed or random mechanism of cellular signalling continues to be debated. However, the role of EVs described in this text, suggests that certain EV populations are released to govern cellular environments. In the context of hematopoietic stem and progenitor cells, EVs act as a key transport mechanism utilized by HSC niche resident cells to instruct HSC functionality.

## References

1 S, C., B, S. & SJ, M. Niches that regulate stem cells and hematopoiesis in adult bone marrow - PubMed. Developmental cell 56 (07/12/2021). 10.1016/j.devcel.2021.05.018

2 Popravko, A., Mackintosh, L. & Dzierzak, E. A life-time of hematopoietic cell function: ascent, stability, and decline. Febs Letters 598 (2024 Mar 5). 10.1002/1873-3468.14843

3 J, C., LSP, H., RJ, d. B. & L, P. Hematopoiesis in numbers - PubMed. Trends in immunology 42 (2021 Dec). 10.1016/j.it.2021.10.006

4 Verovskaya, E. V., Dellorusso, P. V. & Passegué, E. Losing Sense of Self and Surroundings: Hematopoietic Stem Cell Aging and Leukemic Transformation. Trends in molecular medicine 25 (2019 May 17). 10.1016/j.molmed.2019.04.006

5 Ibneeva, L. & Grinenko, T. Frontiers | Dissecting dormancy and quiescence in hematopoietic stem cells. Frontiers in Hematology 3 (2024/09/25). 10.3389/frhem.2024.1401713

6 Chen, Z. et al. Molecular regulation of hematopoietic stem cell quiescence. Cellular and Molecular Life Sciences 2022 79:4 79 (2022-03-31). 10.1007/s00018-022-04200-w

7 Venezia, T. A. et al. Molecular Signatures of Proliferation and Quiescence in Hematopoietic Stem Cells. PLOS Biology 2 (Sep 28, 2004). 10.1371/journal.pbio.0020301

8 Randall, T. D. & Weissman, I. L. Phenotypic and Functional Changes Induced at the Clonal Level in Hematopoietic Stem Cells After 5-Fluorouracil Treatment. Blood 89 (1997/05/15). 10.1182/blood.V89.10.3596

9 Hodgson, G. S., Bradley, T. R., Hodgson, G. S. & Bradley, T. R. Properties of haematopoietic stem cells surviving 5-fluorouracil treatment: evidence for a pre-CFU-S cell? Nature 1979 281:5730 281 (1979-10-01). 10.1038/281381a0

10 J, H.-W. et al. Reduced telomere length variation in healthy oldest old - PubMed. Mechanisms of ageing and development 129 (2008 Nov). 10.1016/j.mad.2008.07.004

11 Niazi, V. et al. The role of genetic/epigenetic factors and microenvironment in hematopoietic stem cell ageing. Biogerontology 2025 26:2 26 (2025-03-22). 10.1007/s10522-025-10218-x

12 Li, X., Wang, J., Hu, L. & Cheng, T. How age affects human hematopoietic stem and progenitor cells and the strategies to mitigate aging. Experimental Hematology 143 (2025/03/01). 10.1016/j.exphem.2025.104711

13 TY, S. et al. Aging is associated with functional and molecular changes in distinct hematopoietic stem cell subsets - PubMed. Nature communications 15 (09/11/2024). 10.1038/s41467-024-52318-1

14 DP, S. Clinical consequences of clonal hematopoiesis of indeterminate potential - PubMed. Blood advances 2 (11/27/2018). 10.1182/bloodadvances.2018020222

15 UM, D. et al. A complex proinflammatory cascade mediates the activation of HSCs upon LPS exposure in vivo - PubMed. Blood advances 6 (06/14/2022). 10.1182/bloodadvances.2021006088

16 S, S., B, J. & JR, K. Protection of hematopoietic stem cells from stress-induced exhaustion and aging - PubMed. Current opinion in hematology 27 (2020 Jul). 10.1097/MOH.0000000000000586

17 Jacobs, K. et al. Stress-triggered hematopoietic stem cell proliferation relies on PrimPol-mediated repriming. Molecular Cell 82 (2022/11/03). 10.1016/j.molcel.2022.09.009

18 Hofmann, J. & Kokkaliaris, K. D. Bone marrow niches for hematopoietic stem cells: life span dynamics and adaptation to acute stress. Blood 144 (2024/07/04). 10.1182/blood.2023023788

19 Busch, C., Nyamondo, K. & Wheadon, H. Complexities of modeling the bone marrow microenvironment to facilitate hematopoietic research. Experimental Hematology 135 (2024/07/01). 10.1016/j.exphem.2024.104233

20 Bacenková, D. et al. Interactions of Hematopoietic and Associated Mesenchymal Stem Cell Populations in the Bone Marrow Microenvironment, In Vivo and In Vitro Model. International Journal of Molecular Sciences 2025, Vol. 26, Page 9036 26 (2025-09-17). 10.3390/ijms26189036

21 Carpenter, R. S., Maryanovich, M., Carpenter, R. S. & Maryanovich, M. Systemic and local regulation of hematopoietic homeostasis in health and disease. Nature Cardiovascular Research 2024 3:6 3 (2024-06-12). 10.1038/s44161-024-00482-4

22 X, Z., C, Z., X, C. & Y, L. Interactions of Hematopoietic Stem Cells with Bone Marrow Niche - PubMed. Methods in molecular biology (Clifton, N.J.) 2346 (2021). 10.1007/7651_2020_298

23 Smith, J. N. P. & Calvi, L. M. Current Concepts in Bone Marrow Microenvironmental Regulation of Hematopoietic Stem and Progenitor Cells. Stem cells (Dayton, Ohio) 31 (2013 Jun). 10.1002/stem.1370

24 Nombela-Arrieta, C. & Manz, M. G. Quantification and three-dimensional microanatomical organization of the bone marrow. Blood Advances 1 (2017/02/14). 10.1182/bloodadvances.2016003194

25 B, C. Normal bone anatomy and physiology - PubMed. Clinical journal of the American Society of Nephrology : CJASN 3 Suppl 3 (2008 Nov). 10.2215/CJN.04151206

26 TJ, V., M, V., GL, N. & LM, M. Multiscale modeling of trabecular bone marrow: understanding the micromechanical environment of mesenchymal stem cells during osteoporosis - PubMed. Journal of biomechanical engineering 137 (2015 Jan). 10.1115/1.4028986

27 F, W., F, M., G, O., L, Z. & S, S. The role of bone marrow on the mechanical properties of trabecular bone: a systematic review - PubMed. Biomedical engineering online 21 (11/23/2022). 10.1186/s12938-022-01051-1

28 Gurevitch, O., Slavin, S. & Feldman, A. G. Conversion of red bone marrow into yellow – Cause and mechanisms. Medical Hypotheses 69 (2007/01/01). 10.1016/j.mehy.2007.01.052

29 Tuljapurkar, S. R. et al. Changes in human bone marrow fat content associated with changes in hematopoietic stem cell numbers and cytokine levels with aging. Journal of Anatomy 219 (2011 Sep 16). 10.1111/j.1469-7580.2011.01423.x

30 J, N., G, F., X, K., J, W. & P, H. Age-related marrow conversion and developing epiphysis in the proximal femur: evaluation with STIR MR imaging - PubMed. Journal of Huazhong University of Science and Technology. Medical sciences = Hua zhong ke ji da xue xue bao. Yi xue Ying De wen ban = Huazhong keji daxue xuebao. Yixue Yingdewen ban 27 (2007 Oct). 10.1007/s11596-007-0537-8

31 M, C., G, R. & C, T. Biogenesis, secretion, and intercellular interactions of exosomes and other extracellular vesicles - PubMed. Annual review of cell and developmental biology 30 (2014). 10.1146/annurev-cellbio-101512-122326

32 Tartaglia, N. R., Martin-Jaular, L., Joliot, A. & Théry, C. Extracellular vesicles: A complex array of particles involved in cell-to-cell communication for tissue homeostasis. Cells & Development (2025/10/24). 10.1016/j.cdev.2025.204054

33 Ripoll, L. et al. Biology and therapeutic potential of extracellular vesicle targeting and uptake. Nature Reviews Molecular Cell Biology 2026 (2026-01-02). 10.1038/s41580-025-00922-4

34 Kwok, Z. H. et al. Extracellular Vesicle Transportation and Uptake by Recipient Cells: A Critical Process to Regulate Human Diseases. Processes 2021, Vol. 9, Page 273 9 (2021-01-31). 10.3390/pr9020273

35 Takeishi, S. et al. Haematopoietic stem cell number is not solely defined by niche availability. Nature 2025 646:8085 646 (2025-08-27). 10.1038/s41586-025-09462-5

36 Feng, C., Fan, H., Tie, R., Xin, S. & Chen, M. Frontiers | Deciphering the evolving niche interactome of human hematopoietic stem cells from ontogeny to aging. Frontiers in Molecular Biosciences 11 (2024/12/04). 10.3389/fmolb.2024.1479605

37 Grenier-Pleau, I. J. et al. Extracellular Vesicles Define Discrete Nano-Based Niches Within the Human Haematopoietic System. Journal of Extracellular Vesicles 14 (2025/12/01). 10.1002/jev2.70181

38 Wells, C. J. et al. Enriching for Extracellular Vesicles from Human Bone. bioRxiv (2025-06-22). 10.1101/2025.06.18.660234

39 Xu, A.-N. et al. Differential Expression of CD49f Discriminates the Independently Emerged Hematopoietic Stem Cells and Erythroid-Biased Progenitors. Blood 134 (2019/11/13). 10.1182/blood-2019-130429

40 Majeti, R., Park, C. Y. & Weissman, I. L. Identification of a Hierarchy of Multipotent Hematopoietic Progenitors in Human Cord Blood. Cell Stem Cell 1 (2007/12/13). 10.1016/j.stem.2007.10.001

41 Chao, M. P., Seita, J. & Weissman, I. L. Establishment of a Normal Hematopoietic and Leukemia Stem Cell Hierarchy. Cold Spring Harbor Symposia on Quantitative Biology 73 (2008-01-01). 10.1101/sqb.2008.73.031

42 Cho, S. H., Chung, I. J., Lee, J. J., Park, M. L. & Kim, H. J. Comparison of CD34+ subsets and clonogenicity in human bone marrow, granulocyte colony-stimulating factor-mobilized peripheral blood, and cord blood. Journal of Korean Medical Science 14 (1999/10/01). 10.3346/jkms.1999.14.5.520

43 Bucar, S. et al. Influence of the mesenchymal stromal cell source on the hematopoietic supportive capacity of umbilical cord blood-derived CD34+-enriched cells. Stem Cell Research & Therapy 2021 12:1 12 (2021-07-13). 10.1186/s13287-021-02474-8

44 Ganan-Gomez, I., Clise-Dwyer, K. & Colla, S. Isolation, culture, and immunophenotypic analysis of bone marrow HSPCs from patients with myelodysplastic syndromes. STAR Protocols 3 (2022/12/16). 10.1016/j.xpro.2022.101764

45 Rodríguez, A. et al. Isolation of human and murine hematopoietic stem cells for DNA damage and DNA repair assays. STAR Protocols 2 (2021 Sep 27). 10.1016/j.xpro.2021.100846

46 Kokkaliaris, K. D. et al. Adult blood stem cell localization reflects the abundance of reported bone marrow niche cell types and their combinations. Blood 136 (2020/11/12). 10.1182/blood.2020006574

47 Lucas, D. Structural organization of the bone marrow and its role in hematopoiesis. Current opinion in hematology 28 (2021 Jan). 10.1097/MOH.0000000000000621

48 Iga, T. et al. Spatial heterogeneity of bone marrow endothelial cells unveils a distinct subtype in the epiphysis. Nature Cell Biology 2023 25:10 25 (2023-10-05). 10.1038/s41556-023-01240-7

49 Spangrude, G. J., Heimfeld, S. & Weissman, I. L. Purification and Characterization of Mouse Hematopoietic Stem Cells. Science 241 (1988-7-1). 10.1126/science.2898810

50 Reya, T. et al. A role for Wnt signalling in self-renewal of haematopoietic stem cells. Nature 2003 423:6938 423 (2003-04-27). 10.1038/nature01593

51 McKenzie, J. L. et al. Individual stem cells with highly variable proliferation and self-renewal properties comprise the human hematopoietic stem cell compartment. Nature Immunology 2006 7:11 7 (2006-10-01). 10.1038/ni1393

52 Jaiswal, S. et al. Age-Related Clonal Hematopoiesis Associated with Adverse Outcomes. New England Journal of Medicine 371 (2014-12-25). 10.1056/NEJMoa1408617

53 Pietras, Eric M. et al. Functionally Distinct Subsets of Lineage-Biased Multipotent Progenitors Control Blood Production in Normal and Regenerative Conditions. Cell Stem Cell 17 (2015/07/02). 10.1016/j.stem.2015.05.003

54 Paul, F. et al. Transcriptional Heterogeneity and Lineage Commitment in Myeloid Progenitors. Cell 163 (2015/12/17). 10.1016/j.cell.2015.11.013

55 Bandyopadhyay, S. et al. Mapping the cellular biogeography of human bone marrow niches using single-cell transcriptomics and proteomic imaging. Cell 187 (2024/06/06). 10.1016/j.cell.2024.04.013

56 Rossi, D. J. et al. Cell intrinsic alterations underlie hematopoietic stem cell aging. Proceedings of the National Academy of Sciences of the United States of America 102 (2005 Jun 20). 10.1073/pnas.0503280102

57 Haan, G. d., Nijhof, W. & Zant, G. V. Mouse Strain-Dependent Changes in Frequency and Proliferation of Hematopoietic Stem Cells During Aging: Correlation Between Lifespan and Cycling Activity. Blood 89 (1997/03/01). 10.1182/blood.V89.5.1543

58 H, G., G, d. H. & MC, F. The ageing haematopoietic stem cell compartment - PubMed. Nature reviews. Immunology 13 (2013 May). 10.1038/nri3433

59 Zhang, P. et al. The physical microenvironment of hematopoietic stem cells and its emerging roles in engineering applications. Stem Cell Research & Therapy 2019 10:1 10 (2019-11-19). 10.1186/s13287-019-1422-7

60 Yang, Z., Dong, R., Mao, X., He, X. C. & Li, L. Stress-protecting harbors for hematopoietic stem cells. Current Opinion in Cell Biology 86 (2024/02/01). 10.1016/j.ceb.2023.102284

61 Mende, N. et al. Unique molecular and functional features of extramedullary hematopoietic stem and progenitor cell reservoirs in humans. Blood 139 (2022/06/09). 10.1182/blood.2021013450

62 Yue, M. et al. Extracellular vesicles remodel tumor environment for cancer immunotherapy. Molecular Cancer 2023 22:1 22 (2023-12-13). 10.1186/s12943-023-01898-5

63 Liu, H. et al. Extracellular vesicles in tumor microenvironment modulation and clinical diagnosis. Pathology - Research and Practice 278 (2026/02/01). 10.1016/j.prp.2025.156339

